# Inhibition of fatty acid amide hydrolase (FAAH) by URB597 counteracts cognitive deficit and alters neuroendocrine stress responses in male and female rats

**DOI:** 10.64898/2025.12.20.695675

**Authors:** Anabela Nagyova, Daniela Jezova, Natasa Hlavacova

## Abstract

Cognitive deficits are hallmark features of several neuropsychiatric disorders, yet therapeutic options remain scarce. Modulation of the endocannabinoid system through inhibition of fatty acid amide hydrolase (FAAH) represents a promising target that may influence both cognitive functions and the neuroendocrine system. However, mechanisms linking FAAH inhibition to these outcomes remain poorly understood. In this study, we hypothesised that FAAH inhibition by URB597 counteracts scopolamine-induced memory deficits and modulates neuroendocrine reactivity differently in males and females. We tested the effects of URB597 (0.3 mg/kg, i.p.) in Sprague Dawley rats at baseline and during a scopolamine challenge (0.5 mg/kg, i.p.). Recognition memory was assessed in the novel object recognition (NOR) task, which also served as a mild stressor. Plasma concentrations of adrenocorticotropic hormone (ACTH), corticosterone, vasopressin, aldosterone, and plasma renin activity (PRA) were measured. URB597 pretreatment counteracted the cognitive impairment induced by scopolamine, showing greater efficacy in males. FAAH inhibition reduced ACTH, corticosterone, vasopressin, and aldosterone concentrations, while PRA remained unaffected. Correlation analyses revealed sex-specific associations. In males, better recognition performance was associated with lower ACTH, corticosterone, and vasopressin, whereas in females, cognition correlated negatively with aldosterone and positively with PRA. These findings demonstrate that FAAH inhibition elicits cognitive protection, associated with the attenuation of neuroendocrine stress responses, and this effect is distinct in males and females. By linking behavioural and endocrine outcomes, this study identifies dual actions of FAAH inhibition and underscores the importance of sex as a biological variable in endocannabinoid-based therapeutic strategies.

## Introduction

Cognitive impairments are common features of psychiatric and neurological disorders and represent a major challenge for effective treatment. Pharmacological interventions that can reverse or attenuate such deficits are therefore of considerable research interest. In this context, the endocannabinoid system has emerged as a promising modulatory system in learning and memory, emotion, and stress responsiveness (Lutz et al., 2015).

The endocannabinoid system comprises endogenous ligands, such as anandamide and 2-arachidonoylglycerol, their metabolic enzymes, and cannabinoid receptors CB1 and CB2. Beyond these canonical targets, endocannabinoids can also act on non-cannabinoid receptors, such as transient receptor potential (TRP) channels, G-protein coupled receptors such as GPR55 and GPR18, and nuclear receptors like PPARs, which broaden their influence on thermoregulation, pain perception, inflammation, and metabolism (Muller et al., 2019; Maccarone et al., 2022). Fatty acid amide hydrolase (FAAH) is a key enzyme responsible for anandamide degradation. Its inhibition increases anandamide levels and enhances endocannabinoid signalling, primarily via CB1 receptor activation (Cravatt et al., 2001; Piomelli, 2003). FAAH inhibitors, e. g. URB597, have been shown to exert anxiolytic, antidepressant-like, and neuroprotective effects in preclinical studies, without the psychotropic side effects typically associated with direct CB1 receptor agonists (Kathuria et al., 2003; Gobbi et al., 2005; Lodola et al., 2015).

Although the behavioural effects of FAAH inhibition have been widely studied, findings regarding its impact on cognition are inconsistent. Several studies have reported cognitive improvements (Varvel et al., 2007; Hlavacova et al., 2015; Rivera et al., 2018), whereas others described neutral or even impairing effects depending on dose, task type, and experimental conditions (Seillier et al., 2010; Basavarajappa et al., 2014; Warren et al., 2022). There are reports that FAAH inhibition may counteract memory deficits in animal models of cognitive dysfunction or memory disturbances (Kruk-Slomka et al., 2019; Oddi et al., 2025). These mixed findings raise the question of whether FAAH inhibition enhances cognition under normal conditions, or whether its benefits emerge primarily under conditions of challenge, such as pharmacologically induced cognitive deficit. Scopolamine, a muscarinic acetylcholine receptor antagonist, can induce such a challenge, as it transiently disrupts cholinergic signalling that is critical for memory formation and recall (Klinkenberg and Blokland, 2010).

The endocannabinoid system plays a critical role in modulating neuroendocrine stress responses. It is known, that activation of CB1 receptors dampens hypothalamic– pituitary–adrenocortical (HPA) axis activity (Patel et al., 2004; Evanson et al., 2010; Hillard, 2018). Acute stress has been shown to increase FAAH activity, resulting in reduced anandamide levels and a concomitant increase in corticosterone concentrations (Hill et al., 2009; Morena et al., 2019). Pharmacological inhibition of FAAH can attenuate this effect, thereby buffering stress-induced increases in corticosterone (Patel et al., 2004). However, a recent systematic review concluded that FAAH inhibitors generally exert little influence on corticosterone under basal conditions, and their capacity to attenuate stress-induced elevations appears to vary across models and experimental conditions (Pereira et al., 2025). Dysregulation of glucocorticoid signalling has long been implicated in cognitive disturbances (De Alcubierre et al. 2023), suggesting that any cognitive effects of FAAH inhibition may in part reflect its impact on glucocorticoid responses. While the main attention has been given to glucocorticoids, evidence linking endocannabinoid signalling to the control of a functionally related hormone vasopressin, is very limited. To our knowledge, there is only one study showing that anandamide can inhibit vasopressin secretion from the posterior pituitary in vitro (Luce et al., 2014). Given the role of vasopressin in HPA axis regulation and memory processes, these sporadic findings indirectly suggest a possible interaction between endocannabinoid signalling and vasopressin, while direct evidence, particularly in the context of FAAH inhibition, is lacking.

The renin–angiotensin–aldosterone system (RAAS) is considered to be another neuroendocrine pathway relevant to stress response and cognition (Correa et al., 2022; Subhani et al., 2025). Plasma renin activity is widely used as an established marker of RAAS function and related sympathetic activation. Within the RAAS, the mineralocorticoid hormone aldosterone represents a key downstream effector. Beyond its classical role in fluid and electrolyte balance, aldosterone has been implicated in stress-related psychiatric disorders (Hlavacova and Jezova, 2008; Hlavacova et al., 2012; Izakova et al., 2023). However, little is known about the effects of endocannabinoid system modulation on RAAS and its potential link to cognition.

Finally, increasing attention has been paid to sex differences in both cognitive processes and the function of the endocannabinoid system. Female rats often display distinct patterns of endocannabinoid tone and stress reactivity compared to males (Reich et al., 2009; Levine et al., 2021). Sex-dependent effects of FAAH inhibitors have been reported, with different behavioural and endocrine outcomes in males and females (Hlavacova et al., 2015; Jankovic et al., 2020). Therefore, it is essential to consider sex as a biological variable when evaluating the effects of endocannabinoid system modulation.

The present study aimed to investigate whether inhibition of FAAH by URB597 can counteract cognitive deficits and alter neuroendocrine stress responses under conditions of acute cholinergic disruption induced by scopolamine. We hypothesised that URB597 would attenuate scopolamine-induced memory deficits in the novel object recognition test. In addition, we expected that URB597 would modulate the neuroendocrine stress response, leading to altered levels of corticosterone, adrenocorticotropic hormone (ACTH), vasopressin, aldosterone and plasma renin activity following behavioural testing. Importantly, we sought to determine whether the cognitive effects of FAAH inhibition would be accompanied by, or potentially reflect, changes in neuroendocrine reactivity. Finally, given known sex differences in both cognitive processing and endocannabinoid system function, we proposed that the effects of URB597 would differ between males and females.

## Material and Methods

### Animals

A total of 40 adult Sprague-Dawley rats, 20 males and 20 females (AnLab, Prague, Czech Republic), weighing 180–230 g and 10 weeks old, were used in this study. Animals were housed two per cage in standard cages with sawdust bedding, with food and water available ad libitum. The housing room was maintained at 22 ± 2 °C with controlled humidity and a reverse 12:12 h light/dark cycle (lights on at 7.00 p.m.). Rats were acclimated to the housing conditions for two weeks before the experiments and were handled daily by the experimenter. All behavioral testing was performed during the dark phase. All procedures were approved by the Animal Health and Animal Welfare Division of the State Veterinary and Food Administration of the Slovak Republic and complied with the NIH Guidelines for the Care and Use of Laboratory Animals.

### Pharmacological treatment

The FAAH inhibitor URB597 (Sigma-Aldrich, Slovakia) was dissolved in dimethyl sulfoxide (DMSO), Tween-80, and 0.9% saline (1:1:8, respectively) and administered intraperitoneally at a dose of 0.3 mg/kg of body weight, as described previously (Hlavacova et al. 2015). Control animals received the vehicle only. To induce cognitive impairment, scopolamine hydrobromide (Sigma-Aldrich, Slovakia) was dissolved in 0.9% saline and administered intraperitoneally at a dose of 0.5 mg/kg. The selected dose of scopolamine was based on previous studies demonstrating reliable induction of cognitive deficits in rats in the novel object recognition test (Hirst et al., 2006; Haider et al., 2016).

### Assessment of cognitive performance

Cognitive performance was assessed using the novel object recognition (NOR) test, as described previously by Hlavacova et al. (2015). One hour before behavioural testing, rats were transported to the experimental room for acclimation. All testing was conducted between 08:30 and 11:30 a.m., during the dark phase of the light/dark cycle. The assessment was repeated twice in the same group of rats, with a one-week washout period between the two experiments. The NOR test was carried out over two consecutive days. On day 1, animals were habituated to the empty arena for 5 min. On day 2, they were exposed to two identical objects during a 15-minute training session. After a 45-minute delay, one familiar object was replaced by a novel one, and animals were allowed to explore for 3 min. Exploration was defined as directing the nose toward or touching the object, while climbing or sitting on it was not considered exploration. Behaviour was video-recorded and analysed using EthoVision XT software (Noldus, Netherlands). Exploration behaviour was analysed by measuring the time spent exploring the familiar object and the time spent exploring the novel object during the test phase. From these measures, two indices were calculated: the recognition index, defined as the proportion of time spent exploring the novel object relative to the total exploration time [recognition index=Tnovel/(Tnovel+Tfamiliar)] and the discrimination index, defined as the difference in exploration times normalised to the total exploration time [discrimination index=(Tnovel−Tfamiliar)/(Tnovel+Tfamiliar).

### Study design

The experimental design combined pharmacological treatment with behavioural testing and subsequent evaluation of neuroendocrine activation. All rats underwent the NOR test on two occasions: first under baseline conditions (Day 2) and again one week later during a scopolamine challenge (Day 9). This repeated-testing design allowed assessment of recognition memory under two conditions, while at the same time, the NOR task served as a mild stressor to elicit neuroendocrine stress response. The experimental design is depicted in Figure 1.

**Figure 1.**
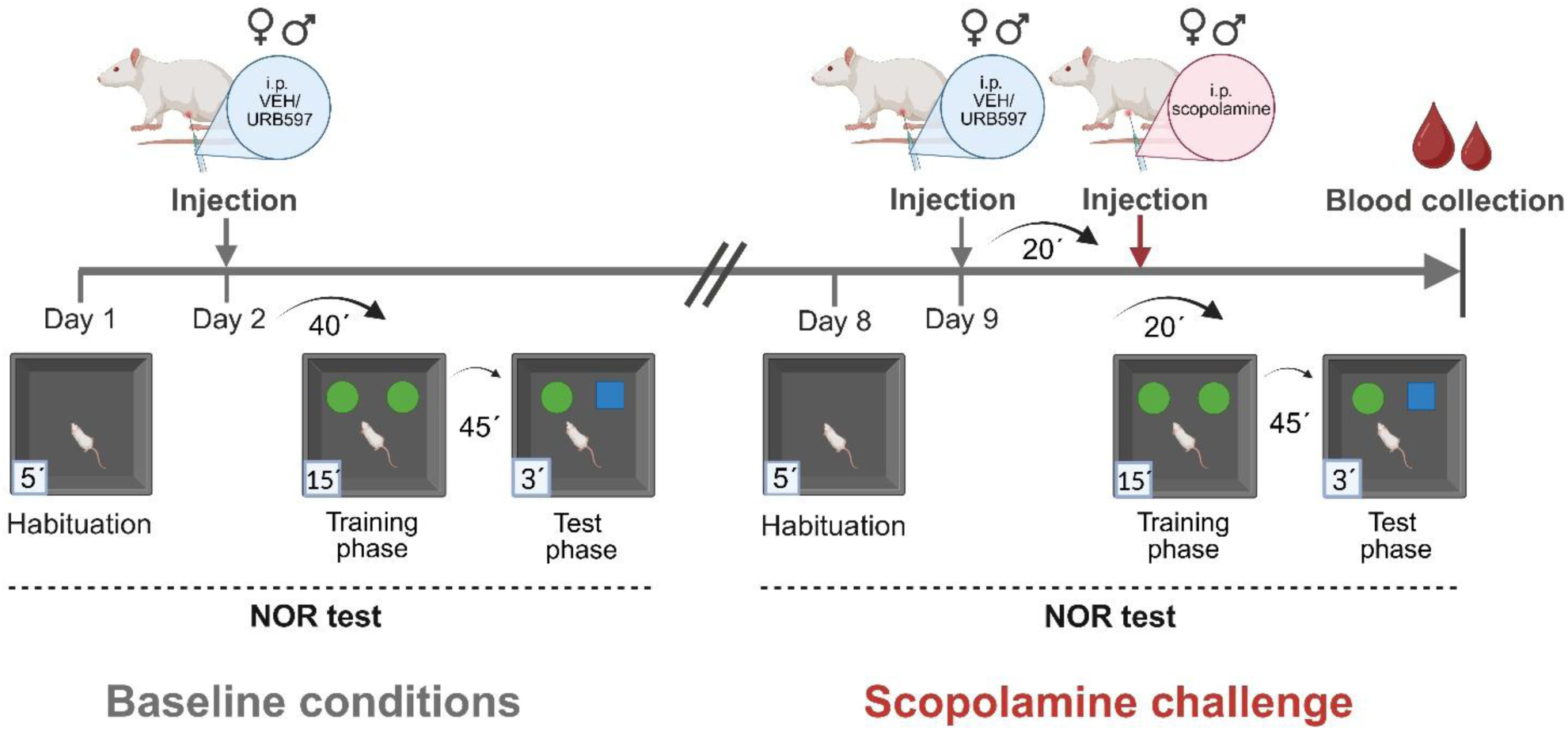
The design of the study. The experimental design combined pharmacological treatment with behavioural testing and subsequent evaluation of neuroendocrine activation. All rats underwent the novel object recognition (NOR) test on two occasions: first under baseline conditions (Day 2) and again one week later during a scopolamine challenge (Day 9). On Day 1, animals were habituated to the arena (5 min). On Day 2, rats received either vehicle (VEH) or URB597 (0.3 mg/kg, i.p.) according to their group assignment, followed 40 min later by the NOR test (training: 15 min; delay: 45 min; test: 3 min). After a 1-week washout, the same cohort was retested. On Day 8, animals underwent a second habituation session. On Day 9, rats were pre-treated with VEH or URB597, followed 20 min later by scopolamine injection (0.5 mg/kg, i.p.) to induce cognitive deficit. The NOR procedure was then repeated, and immediately after the test phase, animals were sacrificed for blood collection to measure endocrine parameters.

Based on sex and treatment, rats were randomly assigned to four experimental groups (n = 10 per group): 1. females-VEH (female rats treated with vehicle), 2. females-URB597 (female rats treated with URB597), 3. males-VEH (male rats treated with vehicle), and 4. males-URB597 (male rats treated with URB597). This group assignment was maintained for the entire experiment.

On Day 1, all animals underwent a 5-minute habituation session of the NOR test without any pharmacological intervention. On Day 2, rats received either URB597 (0.3 mg/kg, i.p.) or vehicle according to their group assignment. Forty minutes after injection, animals were subjected to the training phase of the NOR test, followed by the test phase. After completion of the NOR test, rats were returned to their home cages.

After a one-week wash-out interval, the same cohort of rats was re-tested. On Day 8, animals underwent a second habituation session in the empty arena for 5 min. On Day 9, rats again received URB597 (0.3 mg/kg, i.p.) or vehicle according to their original assignments. Twenty minutes later, all animals were administered scopolamine (0.5 mg/kg, i.p.), and behavioural testing commenced 20 minutes after administration. The NOR procedure was identical to that used on Day 2, consisting of training with two identical objects for 15 min, followed by a 45-minute delay and a 3-minute test phase with one novel object. Immediately after completion of the test phase on Day 9, animals were decapitated to allow the collection of blood for the evaluation of neuroendocrine activation.

### Blood collection and hormone measurements

On Day 9, immediately after completion of the NOR test, all rats were rapidly moved to an adjacent room and sacrificed by decapitation. Trunk blood was collected into cooled polyethylene tubes containing EDTA as an anticoagulant and centrifuged at 4 °C to obtain plasma, which was stored at −20 °C until analysis.

Plasma corticosterone levels were analysed by radioimmunoassay (RIA) after dichloromethane extraction as described earlier (Makatsori et al., 2003). Plasma concentrations of ACTH were determined by RIA described previously (Jezova et al., 1987), using a double antibody technique to separate free and bound fractions. Vasopressin concentrations in plasma were determined by specific RIA, as described previously (Jezova and Michajlovskij, 1992). Plasma aldosterone levels and plasma renin activity were measured by radioimmunoassay (RIA) using commercially available kits (RIA Aldosterone kit, Angiotensin I RIA kit, Immunotech, France). All hormone measurements were performed in duplicate.

### Statistical Analysis

Data were analysed using GraphPad Prism version 10 (GraphPad Software, San Diego, CA). The normality of the data distribution was verified using the Shapiro–Wilk test. Data on NOR test performance under baseline and scopolamine challenge, as well as neuroendocrine data, were analysed by two-way ANOVA with sex and treatment (URB597 vs. vehicle) as between-subject factors. To directly evaluate the effect of scopolamine on recognition memory, repeated-measures ANOVA was performed with condition (baseline vs. scopolamine) as a within-subject factor and treatment and sex as between-subject factors. When significant main effects or interactions were detected, Tukey’s post hoc test was applied. To further explore associations between cognitive performance and neuroendocrine measures, Pearson’s correlation coefficients were calculated between recognition memory indices and concentrations of ACTH, corticosterone, vasopressin, aldosterone and plasma renin activity, separately for males and females. Data are presented as mean ± SEM with individual data points shown. Statistical significance was set at p < 0.05.

## Results

### Cognitive performance in the NOR test at baseline (Day 2)

Under basal conditions, URB597 administration significantly influenced NOR test performance. Two-way ANOVA revealed a significant main effect of treatment on exploration of the familiar object (*F*(1,36) = 6.00, p < 0.05), with URB597-treated animals spending less time with the familiar object compared to vehicle-treated ones (Suppl. Figure 1A). No significant main effect of sex or sex x treatment interaction was found. For novel object exploration, there were significant main effects of sex (*F*(1,36) = 11.29, *p* < 0.01), treatment (*F*(1,36) =15.98, *p* < 0.001), and a sex × treatment interaction (*F*(1,36) =8.49, *p* < 0.01). Post hoc comparisons revealed that URB597 significantly increased novel object exploration in males (*p* < 0.001), while no treatment effect was detected in females (Suppl. Figure 1B).

Analysis of recognition memory indices confirmed these findings. For the recognition index, a significant main effect of treatment (*F*(1,36) = 17.29, *p* < 0.001) and a sex × treatment interaction (*F*(1,36) = 4.32, *p* < 0.05) were observed. The Tukey post hoc test revealed that URB597 increased the recognition index in males compared to vehicle-treated males (*p* < 0.001), whereas no effect was observed in females (Suppl. Figure 1C). Similarly, for the discrimination index, there was a significant main effect of treatment (*F*(1,36) = 17.29, *p* < 0.001) and a sex × treatment interaction (*F*(1,36) = 4.32, *p* < 0.05). URB597-treated males displayed higher discrimination index values compared to vehicle-treated males (*p* < 0.001), while no effect was observed in females (Suppl. Figure 1D).

### Cognitive performance in the NOR test under scopolamine challenge (Day 9)

When rats were retested under a scopolamine challenge, robust effects of URB597 pre-treatment were observed across all measures of recognition memory. For familiar object exploration, two-way ANOVA revealed a significant main effect of treatment (*F*(1,36) = 28.22, *p* < 0.001), with URB597-treated animals spending less time with the familiar object than vehicle controls (Suppl. Figure 2A). For novel object exploration (Suppl. Figure 2B), there was a significant main effect of treatment (*F*(1,36) = 32.29, *p* < 0.001), indicating that URB597 increased novel object exploration compared to vehicle. There was also a significant main effect of sex (*F*(1,36) = 4.23, *p* < 0.05) showing that novel object exploration was higher in males than in females. No significant sex x treatment interaction on object exploration was found.

For the recognition index, there were significant main effects of sex (*F*(1,36) = 4.37, *p* < 0.05) and treatment (*F*(1,36) = 69.17, *p* < 0.001). The recognition index values were higher in males than in females, and URB597 pre-treatment significantly improved the recognition index in both sexes (Suppl. Figure 2C). The discrimination index showed the same pattern. There were significant main effects of sex (*F*(1,36) = 4.37, *p* < 0.05) and treatment (*F*(1,36) = 69.17, *p* < 0.001). Discrimination index values were higher in males and were increased in URB597-pretreated animals compared to vehicle-treated rats (Suppl. Figure 2D).

### Integrated analysis across baseline and scopolamine challenge (Day 2 vs Day 9)

To confirm both the impairing effect of scopolamine and the protective influence of URB597 pre-treatment on cognitive performance in the NOR test, repeated measures ANOVA with condition (baseline vs. scopolamine) as a within-subject factor and treatment and sex as between-subject factors was applied. For familiar object exploration (Figure 2A), a repeated measures ANOVA revealed a significant main effect of treatment (F(1,36) = 22.55, p < 0.001), indicating that URB597-treated rats spent less time exploring the familiar object compared to vehicle-treated rats. There was also a significant main effect of condition (*F*(1,36) = 14.45, *p* < 0.001) and condition × sex interaction (*F*(1,36) = 4.24, *p* < 0.05). Post hoc analysis revealed that URB597-treated males spent significantly less time with the familiar object than vehicle-treated males (*p* < 0.001). URB597-treated females did not differ from vehicle-treated females (*p* < 0.01). These results indicate that the reduction in familiar object exploration was most pronounced in URB597-treated males.

**Figure 2.**
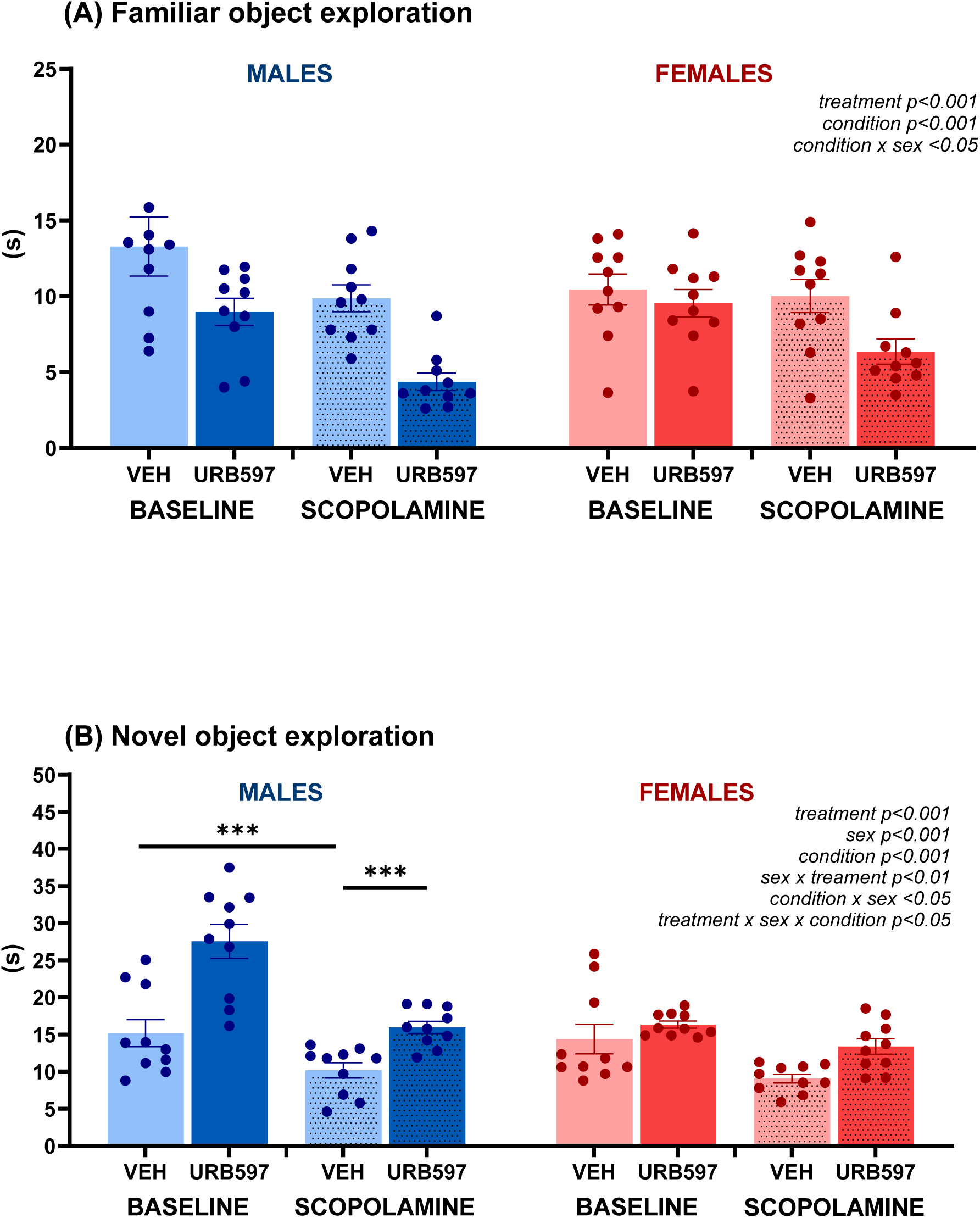
Effects of FAAH inhibition by URB597 on recognition memory in the NOR test under baseline conditions and scopolamine challenge. Time spent exploring familiar (A) and novel (B) objects in male and female rats treated with vehicle (VEH) or URB597 (0.3 mg/kg, i.p.) under baseline conditions and after scopolamine administration (0.5 mg/kg, i.p.). Results are expressed as dot plots, with each dot representing an individual subject and bars indicating mean ± SEM. Statistical analysis as revealed by repeated-measures ANOVA was performed with condition (baseline vs. scopolamine) as a within-subject factor and treatment and sex as between-subject factors. ***p < 0.001.

For novel object exploration (Figure 2B), repeated measures ANOVA revealed robust main effects of treatment (*F*(1,36) = 46.12, *p* < 0.001), sex (*F*(1,36) = 19.07, *p* < 0.001), and condition (*F*(1,36) = 32.52, *p* < 0.001). Rats treated with URB597 spent significantly more time exploring the novel object compared to vehicle-treated controls. Across conditions, males displayed higher exploration times than females. Overall, novel object exploration time was significantly reduced during the scopolamine challenge compared to baseline (*F*(1,36) = 27.58, *p* < 0.001). Importantly, three significant interactions were detected. The sex × treatment interaction (*F*(1,36) = 8.27, p < 0.01) showed that URB597 increased novel object exploration in males (*p* < 0.001), but not in females. The condition × sex interaction (*F*(1,36) = 4.23, *p* < 0.05) indicated that exploration times in females did not differ between baseline and scopolamine challenge, whereas males showed a marked reduction in exploration following scopolamine compared to baseline (*p* < 0.001). Finally, the condition × treatment × sex interaction (*F*(1,36) = 4.79, *p* < 0.05) demonstrated that in vehicle-treated males, scopolamine significantly reduced exploration relative to baseline (*p* < 0.001). This scopolamine-induced impairment was completely reversed by URB597 treatment (URB vs. VEH under scopolamine challenge: *p* < 0.001). In females, URB597 did not significantly alter exploration across conditions. Together, these findings indicate that URB597 enhanced responsiveness to the novel object and partially counteracted scopolamine-induced deficits, with the strongest protective effect observed in males.

For the recognition index (Figure 3A), two-way ANOVA revealed a strong main effect of treatment (*F*(1,36) = 57.58, *p* < 0.001), significant sex × treatment (*F*(1,36) = 5.69, *p* < 0.022) and condition × treatment (*F*(1,36) = 4.71, *p* < 0.05) interactions. Post hoc analysis showed that URB597 markedly improved recognition performance in both females (*p* <0.01) and males (*p* < 0.001) compared with vehicle-treated controls, with the enhancement being greater in males (*p* < 0.05). Furthermore, the treatment × condition interaction indicated that scopolamine impaired recognition performance in vehicle-treated animals (*p* < 0.001), whereas URB597 pretreatment effectively reversed this deficit (*p* < 0.001).

**Figure 3.**
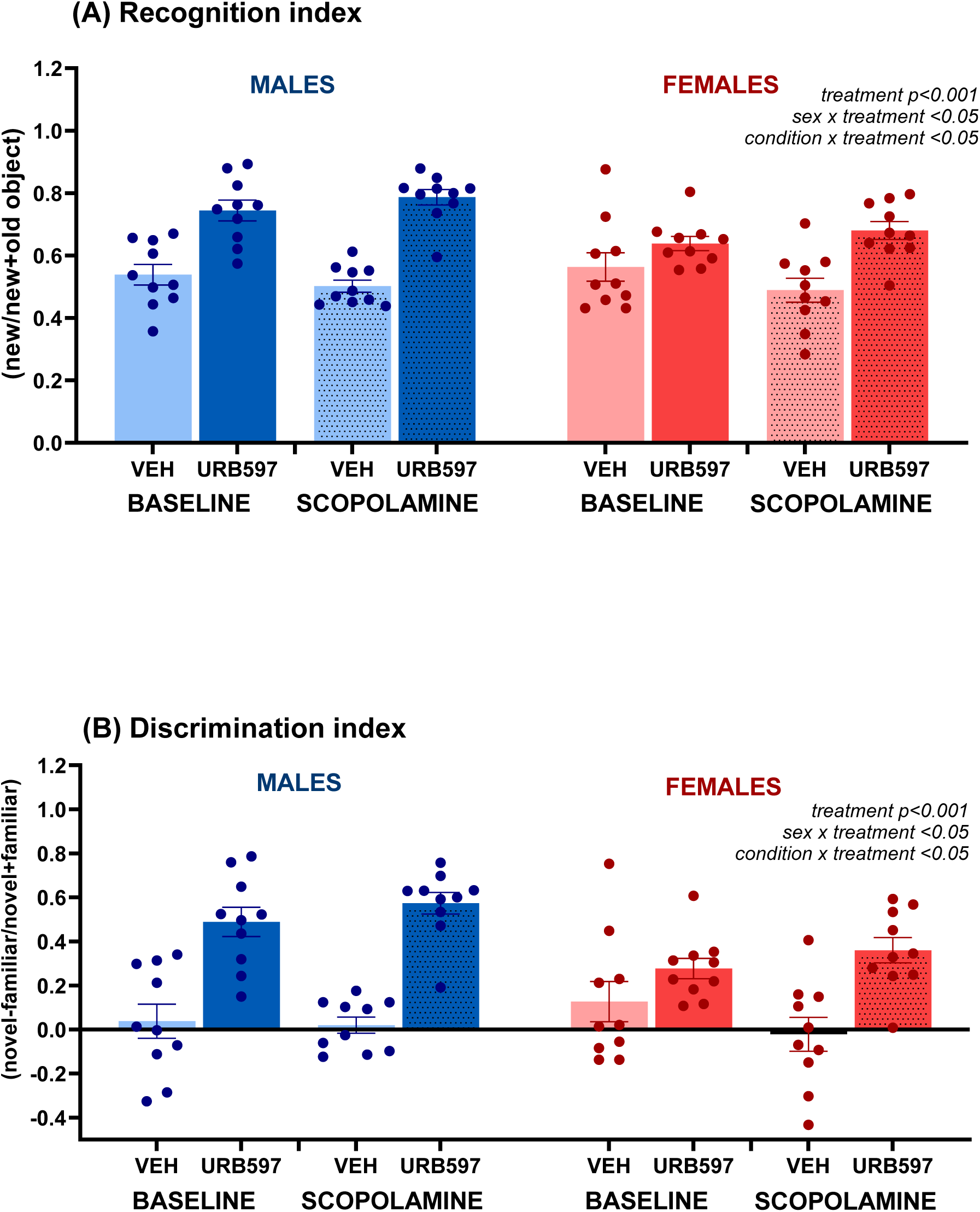
Effects of FAAH inhibition by URB597 on recognition memory in the NOR test during baseline and scopolamine challenge. Recognition index (C), and discrimination index (D) in male and female rats treated with vehicle (VEH) or URB597 (0.3 mg/kg, i.p.) under baseline conditions and after scopolamine administration (0.5 mg/kg, i.p.). Results are expressed as dot plots, with each dot representing an individual subject and bars indicating mean ± SEM. Statistical analysis as revealed by repeated-measures ANOVA was performed with condition (baseline vs. scopolamine) as a within-subject factor and treatment and sex as between-subject factors.

For the discrimination index (Figure 2D), the results closely mirrored those obtained for the recognition index. (Figure 3B). Statistical analysis confirmed a robust main effect of treatment (*F*(1,36) = 57.58, *p* < 0.001), along with significant sex × treatment (*F*(1,36) = 5.69, *p* = 0.022) and condition × treatment (*F*(1,36) = 4.71, *p* = 0.037) interactions. Post hoc comparisons revealed that URB597 significantly improved discrimination performance in both females (*p* < 0.01) and males (*p* < 0.001) compared with vehicle controls, again with a stronger effect in males (*p* < 0.05). The condition × treatment interaction confirmed that URB597 prevented the scopolamine-induced impairment of object discrimination (*p* < 0.001).

### Neuroendocrine parameters

A two-way ANOVA of plasma ACTH concentrations (Figure 4A) revealed a significant main effect of treatment (*F*(1,35) = 14.21, *p* < 0.001), with URB597-treated rats exhibiting lower ACTH levels than vehicle controls, regardless of sex. No significant main effect of sex or interaction between sex and treatment was observed.

**Figure 4.**
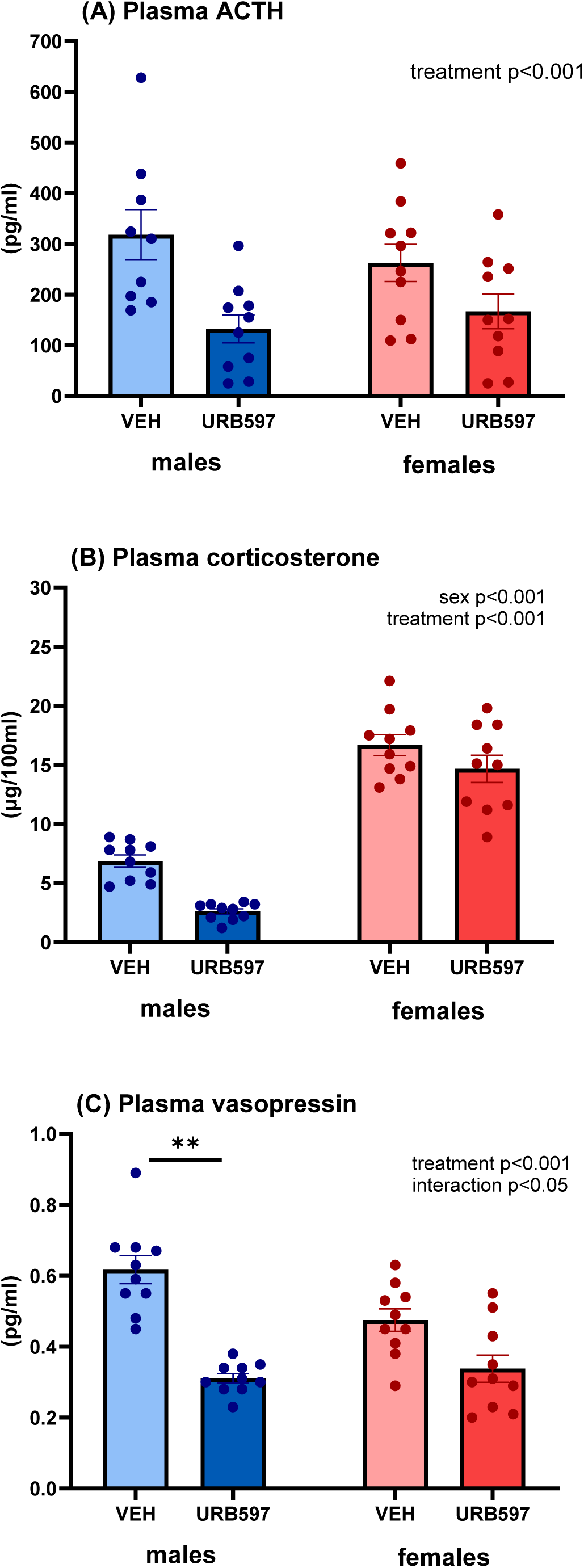
Plasma concentrations of ACTH (A), corticosterone (B), and vasopressin (C) measured after the NOR test in male and female rats treated with vehicle (VEH) or URB597 (0.3 mg/kg). All animals received scopolamine (0.5 mg/kg) before testing. Results are expressed as dot plots, with each dot representing an individual subject and bars indicating mean ± SEM. Statistical analysis as revealed by two-way ANOVA with factors sex and treatment.

For plasma corticosterone concentrations (Figure 4B), two-way ANOVA indicated significant main effects of sex (*F*(1,36) = 198.2, *p* < 0.001) and treatment (*F*(1,36) = 16.39, *p* < 0.001). Females displayed higher corticosterone concentrations than males, and URB597-treated rats had lower levels compared to vehicle-treated rats. No significant sex × treatment interaction was revealed.

Plasma vasopressin concentrations (Figure 4C) were also significantly affected by treatment (*F*(1,36) = 45.93, *p* < 0.001). Treatment with URB597 significantly reduced plasma vasopressin concentrations compared to vehicle. In addition, a significant sex × treatment interaction was found (F(1,36) = 6.69, *p* < 0.05). Post hoc analysis indicated that this effect was primarily driven by males, in which URB597 markedly reduced vasopressin levels compared to vehicle, whereas no significant difference was observed in females.

No significant main effects of sex or treatment, nor a sex × treatment interaction, were found for plasma renin activity (Figure 5A). Plasma aldosterone concentrations (Figure 5B) were significantly influenced by main factors sex (*F*(1,36) = 16.27, *p* < 0.001) and treatment (*F*(1,36) = 4.21, *p* < 0.05). Females showed significantly higher aldosterone concentrations than males. Treatment with URB597 resulted in reduced plasma aldosterone concentrations compared to vehicle. The interaction between factors did not reach statistical significance.

**Figure 5.**
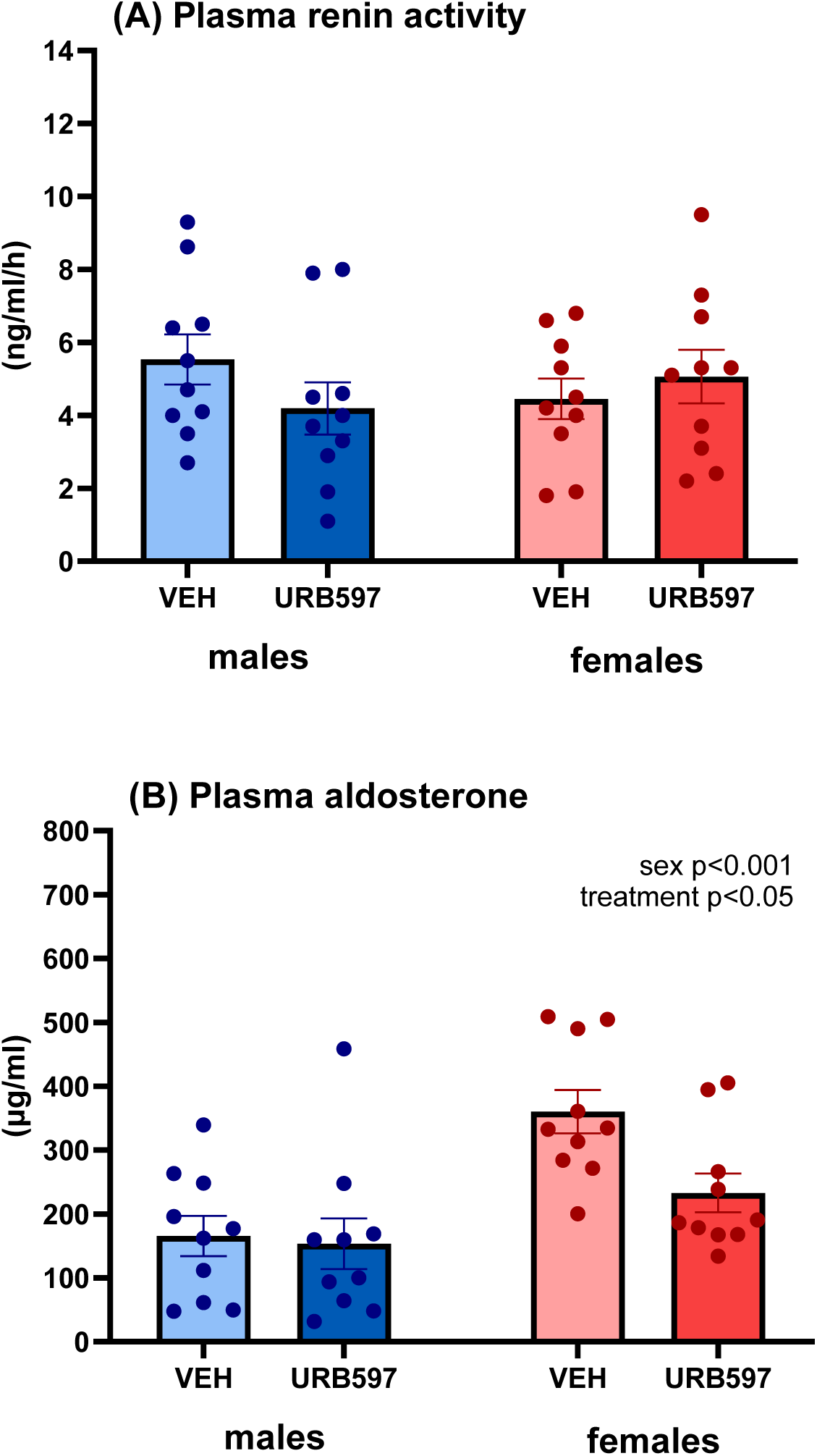
Plasma renin activity (A) and plasma concentrations of aldosterone (B) measured after the NOR test in male and female rats treated with vehicle (VEH) or URB597 (0.3 mg/kg). All animals received scopolamine (0.5 mg/kg) before testing. Results are expressed as dot plots, with each dot representing an individual subject and bars indicating mean ± SEM. Statistical analysis as revealed by two-way ANOVA with factors sex and treatment.

### Correlation analyses

Pearson correlation analyses performed separately for males and females revealed sex-dimorphic patterns of association between cognitive performance and neuroendocrine parameters. In males, both recognition and discrimination indices correlated negatively with concentrations of ACTH (*r* = –0.59; *p* = 0.010), corticosterone (*r* = –0.54; *p* = 0.021), and vasopressin (*r* = –0.73; *p* = 0.001). In females, these associations were not observed; instead, both cognitive indices correlated negatively with aldosterone concentrations (*r* = – 0.67; *p* = 0.002) and positively with plasma renin activity (*r* = 0.53; *p* = 0.025). These findings point to sex-specific differences in the endocrine correlates of recognition memory performance.

## Discussion

In the present study, we have demonstrated that inhibition of FAAH by URB597 improves recognition memory in the NOR task not only under basal conditions, but importantly, also attenuates cognitive deficits induced by the muscarinic antagonist scopolamine. These effects were sex-dependent, with male rats showing the most pronounced cognitive benefits. Moreover, our findings showed that the effects of URB597 involve profound modulation of neuroendocrine stress responses, as evidenced by reduced plasma concentrations of ACTH, corticosterone, vasopressin and aldosterone after the NOR task, which also served as a mild stressor. Importantly, correlation analyses showed that cognitive improvements were accompanied by reductions in endocrine stress markers, with distinct patterns between males and females.

The present data show that FAAH inhibition by URB597 improved recognition memory both at baseline and following scopolamine challenge. These findings are consistent with reports that FAAH inhibition enhances hippocampal plasticity and facilitates memory consolidation under specific conditions (Morena et al., 2014; Rivera et al., 2018; Contarini et al., 2019; Jankovic et al., 2025). Mechanistically, elevated anandamide levels at synapses induced by FAAH inhibition may promote long-term potentiation in the hippocampus and prefrontal cortex, regions critically involved in recognition memory (Colangeli et al., 2017; Lemtiri-Chlieh and Levine, 2022). Furthermore, CB1 receptor activation by increased endocannabinoid tone has been shown to regulate glutamatergic and GABAergic transmission, thereby fine-tuning the excitatory–inhibitory balance required for memory encoding (Busquets-Garcia et al., 2018). Our data add to this framework by showing that FAAH inhibition can specifically buffer the amnestic effects of scopolamine, a widely used pharmacological model of memory impairment (Jagielska et al., 2025). The fact that vehicle-treated rats performed close to chance levels after scopolamine, while URB597-pretreated animals retained preference for the novel object, highlights the robustness of this protective effect and suggests that FAAH inhibition may selectively enhance memory under conditions of cognitive challenge.

Importantly, this memory-enhancing effect was strongly sex-dependent. URB597 markedly improved recognition in males, whereas females showed only modest benefits. Such differences may reflect sex-specific regulation of hippocampal plasticity by the endocannabinoid system, influenced by gonadal hormones, CB1 receptor density, or anandamide turnover (Craft et al., 2013; Hlavacova et al., 2015; Blanton et al., 2021). Recent data further indicate that URB597 modulates neuroplasticity in a sex-dependent manner by suppressing JAK2/STAT3 signalling, enhancing CaMKII activity, and promoting antioxidant responses (Jankovic et al., 2025). Together, these findings suggest that FAAH inhibition enhances cognition most effectively when memory is challenged and that sex-dependent neurobiological mechanisms shape its efficacy.

To the original findings of this work belongs the demonstration that FAAH inhibition modulates neuroendocrine response triggered by behavioural challenge. Previous studies have consistently shown that stress exposure increases FAAH activity, reducing anandamide and thereby facilitating HPA axis activation (Hill et al., 2009; Morena et al., 2019; Maldonado et al., 2020). In our study, FAAH inhibition robustly reduced ACTH and corticosterone concentrations irrespective of sex, confirming its buffering effects on pituitary–adrenocortical activity (Patel et al., 2004; Pereira et al., 2025). However, sex differences emerged in associations between hormonal and cognitive outcomes. Notably, correlation analyses revealed that in males, reductions in ACTH and corticosterone were tightly linked to better cognitive performance, suggesting that FAAH inhibition confers memory benefits partly by buffering HPA axis activation. In contrast, female performance was unrelated to HPA markers.

An intriguing finding is that URB597 reduced plasma vasopressin concentrations, particularly in males. Vasopressin plays a critical role in stress reactivity and social behaviour, and its overactivation has been linked to anxiety, aggression, and stress-related disorders (Neumann & Landgraf, 2012; Carter et al., 2020). Evidence for possible endocannabinoid regulation of vasopressin release is very limited. In vitro studies have shown that anandamide can inhibit vasopressin release from the posterior pituitary via CB1 receptors (Luce et al., 2014), but in vivo endocrine data are lacking. Our results provide the first in vivo evidence that FAAH inhibition decreases vasopressin secretion, extending the role of endocannabinoid signalling beyond HPA axis hormones. The male-specific effect further emphasises the importance of sex differences in endocannabinoid-mediated endocrine regulation. Consistent with this, correlation analyses revealed that in males, lower vasopressin concentrations were associated with better recognition memory, suggesting that suppression of vasopressin release may contribute to the cognitive benefits of FAAH inhibition.

In contrast to the robust effects of URB597 on ACTH, corticosterone, and vasopressin, plasma renin activity remained unaffected by either treatment or sex. This lack of modulation suggests that FAAH inhibition does not uniformly influence all components of the neuroendocrine stress response, but rather acts more selectively. While the RAAS has been implicated in stress and anxiety behavior (Hlavacova and Jezova, 2008; Correa et al., 2022), our data indicate that URB597 did not alter its acute responsiveness under the conditions tested. Interestingly, aldosterone levels in our study did not strictly mirror PRA but instead followed a pattern more closely aligned with ACTH. Females exhibited higher aldosterone concentrations than males, consistent with well-documented sex differences in mineralocorticoid regulation and responsiveness (Viecchola et al. 2024). Importantly, FAAH inhibition by URB597 attenuated aldosterone secretion in both sexes, indicating that endocannabinoid signalling can down-regulate mineralocorticoid output in addition to its established effects on glucocorticoids. Very little is known about the interaction between the endocannabinoid system and the control of aldosterone secretion. Thus, the present data represent a novel finding. Correlation analyses further demonstrated that in females, but not in males, lower aldosterone levels were associated with improved cognitive performance. This sex-specific association suggests that mineralocorticoid signaling may represent a particularly relevant mechanism linking FAAH inhibition to cognitive outcomes in females, in contrast to the HPA axis and vasopressin pathways observed in males.

It should be noted that in the present study, all animals received scopolamine to induce cognitive impairment. In addition to its well-established effects on cholinergic neurotransmission and memory, scopolamine also affects the neuroendocrine system. Muscarinic antagonists have been reported to activate the HPA axis, increasing ACTH and corticosterone secretion, with indications of sex-dependent variation in this response (Smail et al., 2018). Thus, the hormonal levels measured after the NOR test likely reflect not only the stress-inducing properties of the behavioural paradigm but also the pharmacological activation of stress pathways by scopolamine. In this context, the differences observed between URB597- and vehicle-treated animals should be interpreted as treatment-related modulation of a stress response already potentiated by muscarinic antagonism.

Some limitations should be acknowledged. First, the use of scopolamine as a pharmacological model captures only certain aspects of cognitive dysfunction and may not fully reflect the complex pathophysiology of neuropsychiatric disorders. Second, our neuroendocrine measures were obtained at a single time point immediately after behavioural testing, which limits conclusions about the temporal dynamics of stress hormone release. Third, we did not assess endocannabinoid levels directly, so the mechanistic link between FAAH inhibition, endocannabinoid signalling, and HPA axis modulation remains indirect.

## Conclusions

In summary, this study demonstrates that FAAH inhibition with URB597 attenuates scopolamine-induced recognition memory deficits and dampens neuroendocrine stress responses, with pronounced effects in male rats. By simultaneously targeting cognition and HPA axis reactivity, URB597 highlights the potential of endocannabinoid modulation as a therapeutic strategy for stress-related cognitive impairments. The sex-specific patterns observed, with strong cognition–hormone correlations involving the HPA axis and vasopressin in males and a selective aldosterone association in females, underscore the importance of considering sex as a critical biological variable in preclinical and clinical research. These findings provide novel insight into the dual behavioural and endocrine actions of FAAH inhibition and open new avenues for endocannabinoid-based strategies in neuropsychiatric disorders.

## Acknowledgements

This study was supported by the VEGA grant (No. 2/0158/22) from the Scientific Grant Agency of the Ministry of Education, Science, Research and Sport of the Slovak Republic.

---

**Supplementary Figure 1.**
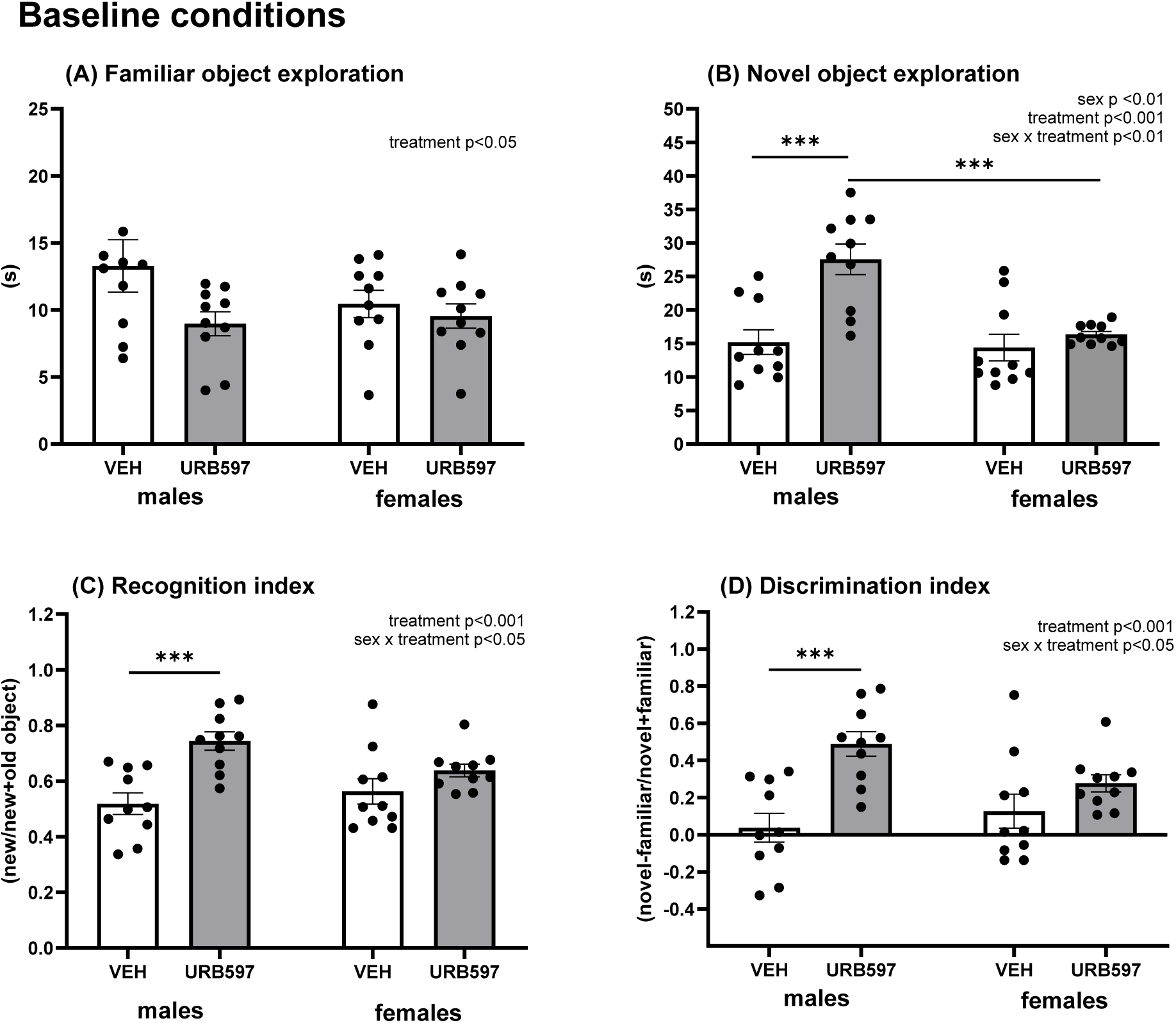
Effects of FAAH inhibition by URB597 on recognition memory in the NOR test under baseline conditions. Time spent exploring familiar (A) and novel (B) objects, recognition index (C), and discrimination index (D) in male and female rats administered with vehicle (VEH) or URB597 (0.3 mg/kg). Results are expressed as dot plots, with each dot representing an individual subject and bars indicating mean ± SEM. Statistical analysis as revealed by twoway ANOVA with factors sex and treatment; ***p < 0.001.

**Supplementary Figure 2.**
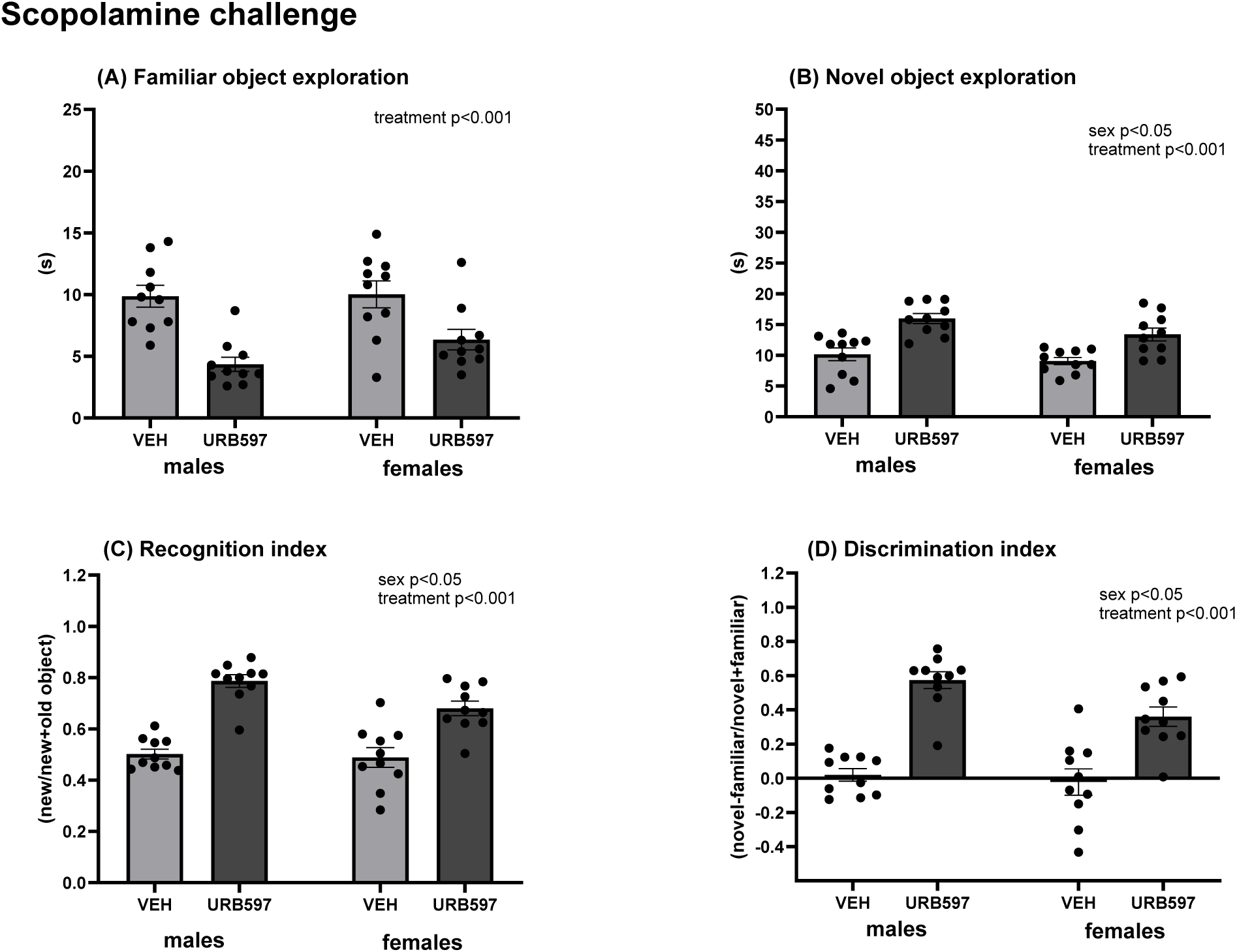
Effects of FAAH inhibition by URB597 on recognition memory in the NOR test under baseline conditions. Time spent exploring familiar (A) and novel (B) objects, recognition index (C), and discrimination index (D) in male and female rats administered with vehicle (VEH) or URB597 (0.3 mg/kg). Results are expressed as dot plots, with each dot representing an individual subject and bars indicating mean ± SEM. Statistical analysis as revealed by twoway ANOVA with factors sex and treatment; ***p < 0.001.

## References

Bangasser, D. A., & Wiersielis, K. R. (2018). Sex differences in stress responses: a critical role for corticotropin-releasing factor. *Hormones (Athens*, Greece), 17(1), 5–13.

Basavarajappa, B. S., Nagre, N. N., Xie, S., & Subbanna, S. (2014). Elevation of endogenous anandamide impairs LTP, learning, and memory through CB1 receptor signaling in mice. Hippocampus, 24(7), 808–818.

Blanton, H. L., Barnes, R. C., McHann, M. C., Bilbrey, J. A., Wilkerson, J. L., & Guindon, J. (2021). Sex differences and the endocannabinoid system in pain. Pharmacology, biochemistry, and behavior, 202, 173107..

Busquets-Garcia, A., Bains, J., & Marsicano, G. (2018). CB1 Receptor Signaling in the Brain: Extracting Specificity from Ubiquity. Neuropsychopharmacology : official publication of the American College of Neuropsychopharmacology, 43(1), 4–20.

Carter C.S., Kenkel W.M., MacLean E.L., et al. (2020). Is oxytocin “nature’s medicine”? Annual Review of Psychology, 71:827–861. doi:10.1146/annurev-psych-010419-050906

Colangeli, R., Pierucci, M., Benigno, A., Campiani, G., Butini, S., & Di Giovanni, G. (2017). The FAAH inhibitor URB597 suppresses hippocampal maximal dentate afterdischarges and restores seizure-induced impairment of short and long-term synaptic plasticity. Scientific reports, 7(1), 11152.

Contarini, G., Ferretti, V., & Papaleo, F. (2019). Acute Administration of URB597 Fatty Acid Amide Hydrolase Inhibitor Prevents Attentional Impairments by Distractors in Adolescent Mice. Frontiers in Pharmacology, 10, 787.

Correa, B. H. M., Becari, L., Fontes, M. A. P., Simões-E-Silva, A. C., & Kangussu, L. M. (2022). Involvement of the Renin-Angiotensin System in Stress: State of the Art and Research Perspectives. Current neuropharmacology, 20(6), 1212–1228.

Cravatt, B. F., Demarest, K., Patricelli, M. P., Bracey, M. H., Giang, D. K., Martin, B. R., & Lichtman, A. H. (2001). Supersensitivity to anandamide and enhanced endogenous cannabinoid signaling in mice lacking fatty acid amide hydrolase. PNAS, 98(16):9371– 9376.

Craft, R. M., Marusich, J. A., & Wiley, J. L. (2013). Sex differences in cannabinoid pharmacology: a reflection of differences in the endocannabinoid system?. Life sciences, 92(8-9), 476–481.

De Alcubierre, D., Ferrari, D., Mauro, G., Isidori, A. M., Tomlinson, J. W., & Pofi, R. (2023). Glucocorticoids and cognitive function: a walkthrough in endogenous and exogenous alterations. Journal of endocrinological investigation, 46(10), 1961–1982.

Evanson, N. K., Tasker, J. G., Hill, M. N., Hillard, C. J., & Herman, J. P. (2010). Fast feedback inhibition of the HPA axis by glucocorticoids is mediated by endocannabinoid signaling. Endocrinology, 151(10), 4811–4819.

Goel, N., Workman, J. L., Lee, T. T., Innala, L., & Viau, V. (2014). Sex differences in the HPA axis. Comprehensive Physiology, 4(3), 1121–1155.

Gobbi, G., Bambico, F. R., Mangieri, R., Bortolato, M., Campolongo, P., Solinas, M., Cassano, T., Morgese, M. G., Debonnel, G., Duranti, A., Tontini, A., Tarzia, G., Mor, M., Trezza, V., Goldberg, S. R., Cuomo, V., & Piomelli, D. (2005). Antidepressant-like activity and modulation of brain monoaminergic transmission by blockade of anandamide hydrolysis. PNAS, 102(51):18620–18625.

Haider, S., Tabassum, S., & Perveen, T. (2016). Scopolamine-induced greater alterations in neurochemical profile and increased oxidative stress demonstrated a better model of dementia: A comparative study. Brain research bulletin, 127, 234–247.

Hill, M. N., McLaughlin, R. J., Morrish, A. C., Viau, V., Floresco, S. B., Hillard, C. J., & Gorzalka, B. B. (2009). Suppression of amygdalar endocannabinoid signaling by stress contributes to activation of the hypothalamic-pituitary-adrenal axis. Neuropsychopharmacology : official publication of the American College of Neuropsychopharmacology, 34(13), 2733–2745.

Hillard C.J. (2018). Circulating endocannabinoids: From whence do they come and where are they going? Neuropsychopharmacology, 43(1):155–172.

Hirst, W. D., Stean, T. O., Rogers, D. C., Sunter, D., Pugh, P., Moss, S. F., Bromidge, S. M., Riley, G., Smith, D. R., Bartlett, S., Heidbreder, C. A., Atkins, A. R., Lacroix, L. P., Dawson, L. A., Foley, A. G., Regan, C. M., & Upton, N. (2006). SB-399885 is a potent, selective 5-HT6 receptor antagonist with cognitive enhancing properties in aged rat water maze and novel object recognition models. European journal of pharmacology, 553(1-3), 109–119.

Hlavacova, N., & Jezova, D. (2008). Chronic treatment with the mineralocorticoid hormone aldosterone results in increased anxiety-like behavior. Hormones and behavior, 54(1), 90–97.

Hlavacova N., Chmelova M., Danevova V., Csanova A., Jezova D. (2015). Inhibition of fatty-acid amide hydrolase (FAAH) exerts cognitive improvements in male but not female rats. Endocrine Regulations, 49(3):131–136.

Jankovic, M., Spasojevic, N., Ferizovic, H., Stefanovic, B., & Dronjak, S. (2020). Inhibition of the fatty acid amide hydrolase changes behaviors and brain catecholamines in a sex-specific manner in rats exposed to chronic unpredictable stress. Physiology & behavior, 227, 113174.

Jankovic, M., Spasojevic, N., Ferizovic, H., Stefanovic, B., Virijevic, K., & Dronjak, S. (2025). URB597 modulates neuroplasticity, neuroinflammatory, and Nrf2/HO-1 signaling pathways in the hippocampus and prefrontal cortex of male and female rats in a stress-induced model of depression. Physiology & behavior, 295, 114893.

Jezova, D., Kvetnanský, R., Kovács, K., Oprsalová, Z., Vigas, M., & Makara, G. B. (1987). Insulin-induced hypoglycemia activates the release of adrenocorticotropin predominantly via central and propranolol insensitive mechanisms. Endocrinology, 120(1), 409–415.

Jezova, D., & Michajlovskij, N. (1992). N-methyl-D-aspartic acid injected peripherally stimulates oxytocin and vasopressin release. Endocrine regulations, 26(2), 73–75.

Kathuria, S., Gaetani, S., Fegley, D., Valiño, F., Duranti, A., Tontini, A., Mor, M., Tarzia, G., La Rana, G., Calignano, A., Giustino, A., Tattoli, M., Palmery, M., Cuomo, V., & Piomelli, D. (2003). Modulation of anxiety through blockade of anandamide hydrolysis. Nature Medicine, 9(1):76–81.

Klinkenberg, I., & Blokland, A. (2010). The validity of scopolamine as a pharmacological model for cognitive impairment: a review of animal behavioral studies. Neuroscience and biobehavioral reviews, 34(8), 1307–1350.

Kruk-Slomka, M., Banaszkiewicz, I., Slomka, T., & Biala, G. (2019). Effects of Fatty Acid Amide Hydrolase Inhibitors Acute Administration on the Positive and Cognitive Symptoms of Schizophrenia in Mice. Molecular neurobiology, 56(11), 7251–7266.

Lemtiri-Chlieh, F., & Levine, E. S. (2022). 2-AG and anandamide enhance hippocampal long-term potentiation *via* suppression of inhibition. Frontiers in cellular neuroscience, 16, 1023541.

Levine, A., Liktor-Busa, E., Lipinski, A. A., Couture, S., Balasubramanian, S., Aicher, S. A., Langlais, P. R., Vanderah, T. W., & Largent-Milnes, T. M. (2021). Sex differences in the expression of the endocannabinoid system within V1M cortex and PAG of Sprague Dawley rats. Biology of sex differences, 12(1), 60.

Lodola A., Mulholland K., Rivara S., & Rivara, S. (2015). Fatty acid amide hydrolase inhibitors: A patent review (2009–2014). Expert Opinion on Therapeutic Patents, 25(10):1247–1276.

Luce, V., Fernandez Solari, J., Rettori, V., & De Laurentiis, A. (2014). The inhibitory effect of anandamide on oxytocin and vasopressin secretion from neurohypophysis is mediated by nitric oxide. Regulatory peptides, 188, 31–39.

Lutz B., Marsicano G., Maldonado R., Hillard C.J. (2015). The endocannabinoid system in guarding against fear, anxiety and stress. Nature Reviews Neuroscience, 16(12):705–718. doi:10.1038/nrn4036

Maccarrone, M., Di Marzo, V., Gertsch, J., Grether, U., Howlett, A. C., Hua, T., Makriyannis, A., Piomelli, D., Ueda, N., & van der Stelt, M. (2023). Goods and Bads of the Endocannabinoid System as a Therapeutic Target: Lessons Learned after 30 Years. Pharmacological reviews, 75(5), 885–958.

Maldonado, R., Cabañero, D., & Martín-García, E. (2020). The endocannabinoid system in modulating fear, anxiety, and stress . Dialogues in clinical neuroscience, 22(3), 229–239.

Makatsori, A., Duncko, R., Schwendt, M., Moncek, F., Johansson, B. B., & Jezova, D. (2003). Voluntary wheel running modulates glutamate receptor subunit gene expression and stress hormone release in Lewis rats. Psychoneuroendocrinology, 28(5), 702–714.

Morena, M., Roozendaal, B., Trezza, V., Ratano, P., Peloso, A., Hauer, D., Atsak, P., Trabace, L., Cuomo, V., McGaugh, J. L., Schelling, G., & Campolongo, P. (2014). Endogenous cannabinoid release within prefrontal-limbic pathways affects memory consolidation of emotional training. Proceedings of the National Academy of Sciences of the United States of America, 111(51), 18333–18338.

Morena, M., Aukema, R. J., Leitl, K. D., Rashid, A. J., Vecchiarelli, H. A., Josselyn, S. A., & Hill, M. N. (2019). Upregulation of Anandamide Hydrolysis in the Basolateral Complex of Amygdala Reduces Fear Memory Expression and Indices of Stress and Anxiety. The Journal of neuroscience : the official journal of the Society for Neuroscience, 39(7), 1275–1292.

Muller, C., Morales, P., & Reggio, P. H. (2019). Cannabinoid Ligands Targeting TRP Channels. Frontiers in molecular neuroscience, 11, 487.

Neumann I.D., Landgraf R. (2012). Balance of brain oxytocin and vasopressin: Implications for anxiety, depression, and social behaviors. Trends in Neurosciences, 35(11):649–659.

Oddi, S., Scipioni, L., Totaro, A., Giacovazzo, G., Ciaramellano, F., Tortolani, D., Leuti, A., Businaro, R., Armeli, F., Bilkei-Gorzo, A., Coccurello, R., Zimmer, A., & Maccarrone, M. (2025). Fatty-acid amide hydrolase inhibition mitigates Alzheimer’s disease progression in mouse models of amyloidosis. The FEBS journal, 292(16), 4160–4182.

Patel, S., Roelke, C. T., Rademacher, D. J., Cullinan, W. E., & Hillard, C. J. (2004). Endocannabinoid signaling negatively modulates stress-induced activation of the hypothalamic-pituitary-adrenal axis. Endocrinology, 145(12), 5431–5438.

Pereira, C. F., Boileau, I., & Kloiber, S. (2025). Effects of pharmacological inhibition of fatty acid amide hydrolase on corticosterone release: a systematic review of preclinical studies. Discover mental health, 5(1), 51.

Piomelli D. (2003). The molecular logic of endocannabinoid signalling. Nature Reviews Neuroscience, 4(11):873–884.

Portugalov, A., & Akirav, I. (2024). FAAH Inhibition Reverses Depressive-like Behavior and Sex-Specific Neuroinflammatory Alterations Induced by Early Life Stress. Cells, 13(22), 1881.

Reich C.G., Taylor M.E., McCarthy M.M. (2009). Differential effects of chronic unpredictable stress on hippocampal CB1 receptors in male and female rats. Neuroscience, 158(2):523–530. doi:10.1016/j.neuroscience.2008.10.041

Rivera, P., Fernández-Arjona, M. D. M., Silva-Peña, D., Blanco, E., Vargas, A., López-Ávalos, M. D., Grondona, J. M., Serrano, A., Pavón, F. J., Rodríguez de Fonseca, F., & Suárez, J. (2018). Pharmacological blockade of fatty acid amide hydrolase (FAAH) by URB597 improves memory and changes the phenotype of hippocampal microglia despite ethanol exposure. Biochemical pharmacology, 157, 244–257.

Seillier, A., Advani, T., Cassano, T., Hensler, J. G., & Giuffrida, A. (2010). Inhibition of fatty-acid amide hydrolase and CB1 receptor antagonism differentially affect behavioural responses in normal and PCP-treated rats. The international journal of neuropsychopharmacology, 13(3), 373–386.

Smail, M. A., Soles, J. L., Karwoski, T. E., Rubin, R. T., & Rhodes, M. E. (2018). Sexually diergic hypothalamic-pituitary-adrenal axis responses to selective and non-selective muscarinic antagonists prior to cholinergic stimulation by physostigmine in rats. Brain research bulletin, 137, 23–34

Subhani, M. K., Sabani, A., Omer, O., Kheirelsid, F., & Srour, A. K. (2025). Impact of Renin-Angiotensin-Aldosterone System Modulation on Cognitive and Neuropsychiatric Outcomes: A Systematic Review of Clinical and Mechanistic Evidence. Cureus, 17(7), e87557.

Steckelings U.M., et al. (2012). The renin–angiotensin system in the brain: Its role in cardiovascular, stress and behavior. Pharmacological Research, 65(6):640–652. doi:10.1016/j.phrs.2012.03.003

Varvel, S. A., Wise, L. E., Niyuhire, F., Cravatt, B. F., & Lichtman, A. H. (2007). Inhibition of fatty-acid amide hydrolase accelerates acquisition and extinction rates in a spatial memory task. Neuropsychopharmacology 32(5), 1032–1041.

Viecchiola, A., Uslar, T., Friedrich, I., Aguirre, J., Sandoval, A., Carvajal, C. A., Tapia-Castillo, A., Martínez-García, A., & Fardella, C. E. (2024). The role of sex hormones in aldosterone biosynthesis and their potential impact on its mineralocorticoid receptor. Cardiovascular endocrinology & metabolism, 13(3), e0305.

Warren, W. G., Papagianni, E. P., Hale, E., Brociek, R. A., Cassaday, H. J., & Stevenson, C. W. (2022). Endocannabinoid metabolism inhibition has no effect on spontaneous fear recovery or extinction resistance in Lister hooded rats. Frontiers in Pharmacology, 13, 1082760.

